# Fixation-locked hippocampal activity reflects semantic content and temporal order of visual exploration during scene encoding

**DOI:** 10.64898/2026.05.15.725376

**Authors:** Arantzazu San Agustín, Joel L. Voss, James E. Kragel

## Abstract

Memory formation relies on the hippocampus and unfolds over time across experience, such as during the visual exploration of complex, naturalistic scenes. Eye movements evoke hippocampal activity, including fixation-locked field potentials and phase resets of theta oscillations. This suggests that hippocampal encoding is temporally structured by the sequence of visual fixations. Because eye-movement sequences sample semantically meaningful portions of scenes, they provide temporal structure to semantic content in memory. However, it remains unclear how the semantic content and temporal order of fixations jointly shape medial temporal lobe activity. We therefore tested whether intracranial EEG recordings from human hippocampus and amygdala reflect the semantic content and temporal order of individual fixations during encoding of naturalistic scenes. Relative to other semantic content, fixations on people were particularly relevant for memory, with the first fixation on a person predicting subsequent scene recognition. Fixation-locked hippocampal responses were enhanced for fixations to people relative to other semantic content, expressed in both larger fixation-evoked potentials and stronger theta phase locking. These effects were strongest for the first fixation relative to subsequent fixations. Theta phase locking was also enhanced in both hippocampus and amygdala for first fixations on people relative to later fixations and to other semantic content. These findings indicate that medial temporal lobe activity is structured by discrete fixation-level events during scene encoding, suggesting that theta-paced sampling contributes to the transformation of semantic and temporal components of visual experiences into memory.

**Significance Statement:** This study shows that the semantic content and order of eye fixations jointly influence human hippocampal activity during memory encoding. Combining intracranial recordings, eye-movement tracking, and deconvolutional modeling, we show that the first glance at a person within naturalistic scenes is a privileged event, associated with increased hippocampal activity, theta-phase resetting in hippocampus and amygdala, and subsequent memory success. These findings recast eye movements not as mere motor acts, but as an important process that helps medial-temporal structures prioritize and integrate behaviorally relevant information into episodic memory.

## Introduction

Episodic memory reflects the binding of what was experienced (semantic content) with when and where it occurred (spatiotemporal context; Tulving, 2002; Eichenbaum, 2017). In primates, visual scenes are sampled via saccadic eye movements, which discretize semantic content from individual fixations into a spatiotemporal sequence (Liversedge et al., 2011). Thus, eye movements may structure visual experience into the sequences that form episodic memories (Henderson et al., 2005; Wynn et al., 2016; Kragel and Voss, 2021). Although hippocampus and amygdala are critical for episodic and semantic memory (Scoville and Milner, 1957; Cahill et al., 1995; Manns et al., 2003; Phelps, 2004),relatively little work has examined whether their mnemonic functions are organized by eye movements (Voss et al., 2017).

The hippocampus supports semantic discrimination by encoding relational structure (Eichenbaum, 2004) through neuronal populations sensitive to semantic content (Kreiman et al., 2000; Quiroga et al., 2005; Quiroga, 2012) and spatiotemporal context (Umbach et al., 2020; Mackay et al., 2024; Bausch et al., 2026). This relational processing also extends to visual exploration. Hippocampal activity predicts memory expression in eye movements (Hannula and Ranganath, 2009), and fixation-locked theta phase resets in hippocampus are linked to memory formation (Jutras et al., 2013; Kragel et al., 2020). Theta oscillations in particular are closely associated with episodic memory formation (Lega et al., 2012; Cruzat et al., 2021), and their phase locking at fixation onset suggests hippocampal encoding is temporally structured by eye movements (Hoffman et al., 2013; Jutras et al., 2013).

The amygdala, a heterogeneous structure comprising functionally distinct nuclei (Janak and Tye, 2015), signals the behavioral relevance of stimuli across a broad range of categories, influencing both memory consolidation and attention toward significant stimuli (Adolphs, 2010). This contribution to visual exploration varies with the nature of behavioral relevance. The amygdala is involved in (Minxha et al., 2017; Lee et al., 2026) and even necessary for (Kaskan et al., 2022) orienting to stimuli with innate biological value such as conspecifics. Further, its neurons integrate motivational significance with spatial information to guide attention (Peck et al., 2013; Peck and Salzman, 2014). However, this prior work has not isolated scene-level and fixation-level contributions of these structures, leaving their mnemonic and attentional functions unclear.

Beyond which brain structures are engaged, the order (Damiano and Walther, 2019) and semantic content (Foulsham and Underwood, 2008; Lyu et al., 2020; Broers et al., 2022) of fixations jointly shape what is remembered. Early fixations are driven toward semantically discrepant objects (Loftus and Mackworth, 1978; De Graef et al., 1990) as well as faces and people, which elicit earlier and longer fixations with better memory (Guo et al., 2006; Fletcher-Watson et al., 2008). Scanpath similarity between study and test correlates with increased memory, especially for the earliest fixations within the scanpath (Wynn et al., 2016). Early fixations may therefore support memory by providing a temporal reference frame for subsequently encoded content. Despite this evidence, memory research typically treats visual input as static, without measuring or modeling gaze behavior, thereby limiting inferences about dynamic processes organized by fixations (Kragel and Voss, 2022).

Motivated by these considerations, we combined the high spatiotemporal resolution of intracranial electroencephalography (iEEG) recordings from the human hippocampus and amygdala with eye tracking to test whether neural activity is modulated by the semantic content and temporal order of individual fixations during scene encoding. A key challenge was that visual information, fixations, and saccades can produce overlapping neural signals (Ringo et al., 1994; Hoffman et al., 2013; Doucet et al., 2020; Katz et al., 2020), making it difficult to isolate fixation-specific activity. To overcome this issue, we applied a regression-based deconvolution approach (Ehinger and Dimigen, 2019) to isolate neural activity related to certain fixations. Complementing this time-domain analysis, we characterized how semantic content and temporal order modulate theta phase locking to fixations. Together, these approaches allowed us to reveal the fixation-level neural processes linking the semantic content and temporal order of visual exploration to memory formation.

## Material and methods

### Participants

We analyzed data from six participants (three female; mean age = 29 years, range = 24 − 38) with medically refractory epilepsy collected at the Northwestern Memorial Hospital Comprehensive Epilepsy Center (Chicago, IL). All participants had depth electrodes implanted as part of clinical monitoring and localization of epileptogenic tissue prior to surgical treatment. Written informed consent was acquired prior to data collection in accordance with the Northwestern University Institutional Review Board. Analyses of data from these participants have been reported (Kragel et al., 2021), with no overlap of current with previously reported analyses.

### Recognition Memory Task

Participants performed a recognition memory task that comprised eight study-test blocks. In each block, they studied a sequence of 24 naturalistic scenes followed by a recognition test for 24 novel and 24 repeated scenes, presented in randomized order. Stimuli were selected from the Microsoft COCO dataset and depicted everyday scenes containing common objects in natural contexts (Lin et al., 2014). Each studied list comprised scenes from three semantic categories: People, Animals, and Food (8 scenes of each category per block). Novel scenes were matched for content to the repeated scenes (Figure 1A).

**Figure 1.**
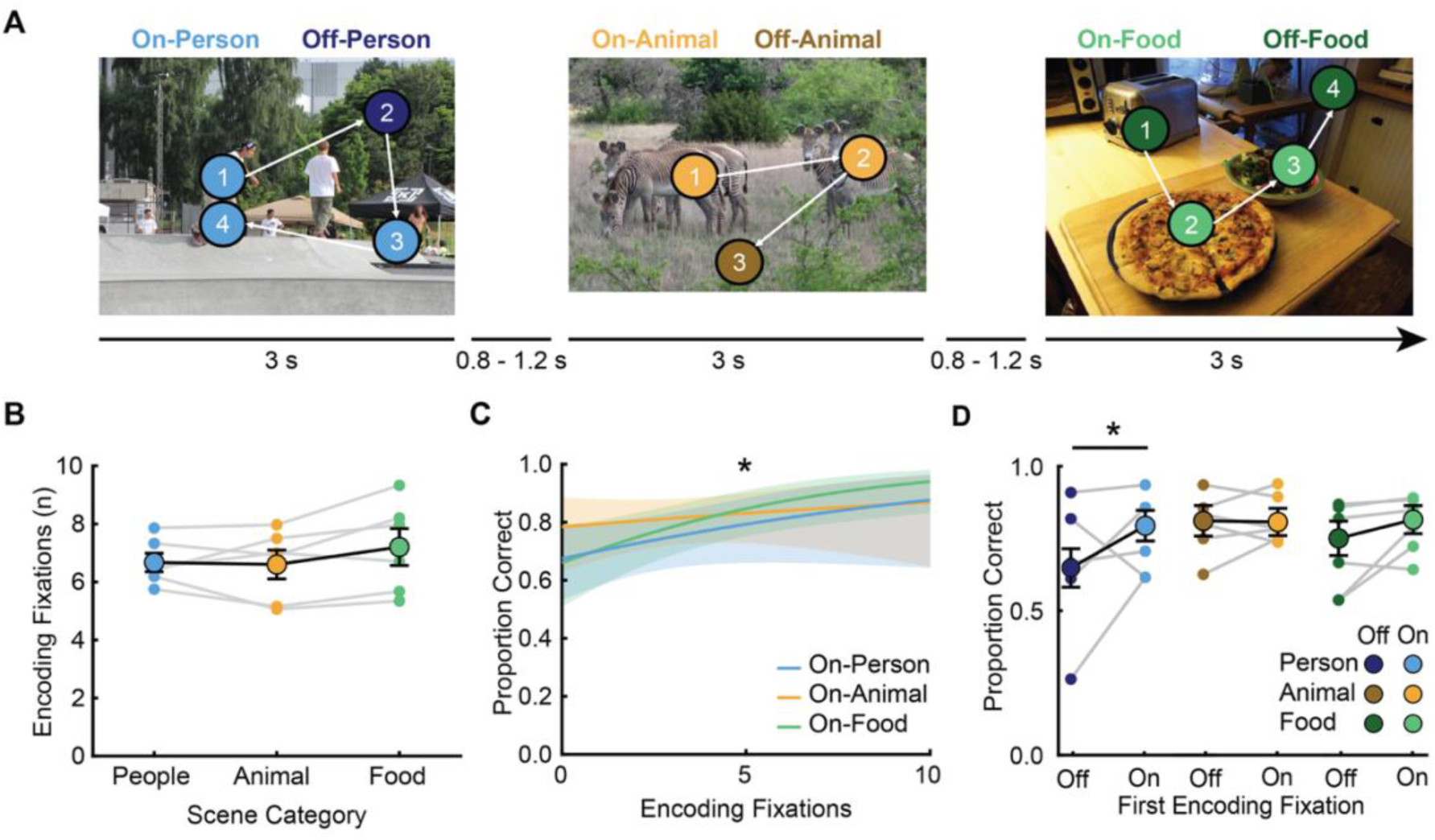
Semantic content and temporal order of fixations during encoding predict subsequent memory. **A.** Representative scenes and timing during the study phase of the task for the 3 different scene semantic categories: People, Animals, and Food. Examples of fixation sequences are depicted showing fixation conditions and order (blue for People, yellow for Animals, green for Food; light colors to On-Category fixations, dark colors to Off-Category fixations). **B.** Number of fixations did not differ significantly across categories. Large points and error bars denote the group mean and standard error. Small points denote the average number of fixations for each participant. **C.** Number of On-Category fixations per scene predicted subsequent recognition accuracy. Lines indicate predicted effects from linear-mixed models. Shaded regions denote SEM across participants. **D.** Semantic content of the first fixation predicted recognition accuracy selectively for People scenes. Recognition accuracy at test for People scenes was significantly greater with On-Person compared to Off-Person first fixations. * *p < 0.05*.

During the study phase, scenes were displayed for 3 seconds, followed by a randomly jittered inter-trial interval of 0.8 to 1.2 seconds. The test phase of the task used a gaze-contingent design to allow memory to guide visual sampling (see Kragel et al. 2021 for more detailed description). Test trials were displayed until the participant indicated by button press if the stimulus was repeated or novel (old/new). Test trials were followed by a 0.8 to 1.2 second inter-trial interval. Prior to each study and test trial, a centrally located fixation cross was presented for 0.5 seconds to alert the participant about the upcoming trial. Note that the test phase was used here only to categorize the success of encoding based on subsequent memory performance, with all current analyses of eye movements and iEEG data limited to the study phase.

### Eye Tracking and Fixation Categories

Eye movements were recorded at 500 Hz using an EyeLink 1000 tracking system (SR Research, Ontario, Canada). The camera was positioned beneath the display monitor, which was located approximately 60 cm from the participant’s eyes. Eye-tracking calibration was conducted before the study phase of each block using a five-point calibration procedure, which was repeated if the mean error exceeded 1.5° of visual angle. Blinks, saccades, and fixations were detected from the continuous eye-movement records. Blinks were identified by pupil size. Saccades were identified by thresholds for motion (0.15°), velocity (30°/s), and acceleration (8000°/s^2^), and the remaining epochs below detection thresholds were categorized as fixations. The average gaze position during the fixation period was used to calculate the location of each fixation event.

Pixel-wise annotations and per-instance segmentation masks of COCO images were used to assign semantic categories to fixations based on their spatial correspondence. For each fixation, its screen coordinates (x, y) were mapped onto the image space and tested for overlap with object segmentation masks. A given semantic category was assigned to a fixation when it fell within the pixel boundaries of an annotated object instance. Fixations landing on People, Animals, or Food regions were assigned accordingly and labeled as On-Category fixations (On-Person, On-Animal, or On-Food). Fixations falling on other areas of the scene were labeled as Off-Category fixations (Off-Person, Off-Animal, or Off-Food; Figure 1A).

Fixation order was defined relative to scene onset. The first fixation was defined as the first fixation that began after scene presentation. Participants were initially fixating a central cross; therefore, any fixation already ongoing at scene onset (i.e., starting during the fixation-cross period) was excluded. Subsequent fixations were then indexed sequentially based on their onset times.

### iEEG recordings and electrode localization

Stereotactic depth electrodes (contacts spaced 5 to 10 mm apart, AD-TECH Medical Instrument Co., Racine, WI) were implanted for clinical purposes and provided coverage of the hippocampus and amygdala. Electrode localization was performed by co-registering presurgical T1-weighted structural MRIs and postimplant computed tomography (CT) images using SPM12 (Ashburner and Friston, 1997). T1-weighted images were normalized to MNI (Montreal Neurological Institute) space by using a combination of affine and nonlinear registration steps, bias correction, and tissue segmentation into gray matter, white matter, and cerebrospinal fluid components (Gaser et al., 2024). The resulting deformation fields were applied to individual electrode locations identified on postimplant CT images using Bioimage Suite (Papademetris et al., 2006). The classification of electrodes within hippocampus (Figure 2A) and amygdala (Figure 2B) was conducted first by an automated comparison of the specific MNI coordinates of each electrode contact with the AAL3 atlas parcellation of left and right hippocampus and amygdala and verified after by visual inspection of co-registered native space scans. Only electrodes confirmed to be located within hippocampus (n = 25; mean = 4.2 per participant, range 2 – 7) and amygdala (n = 24; mean = 4.0 per participant, range 3 – 6) by visual inspection were retained.

**Figure 2.**
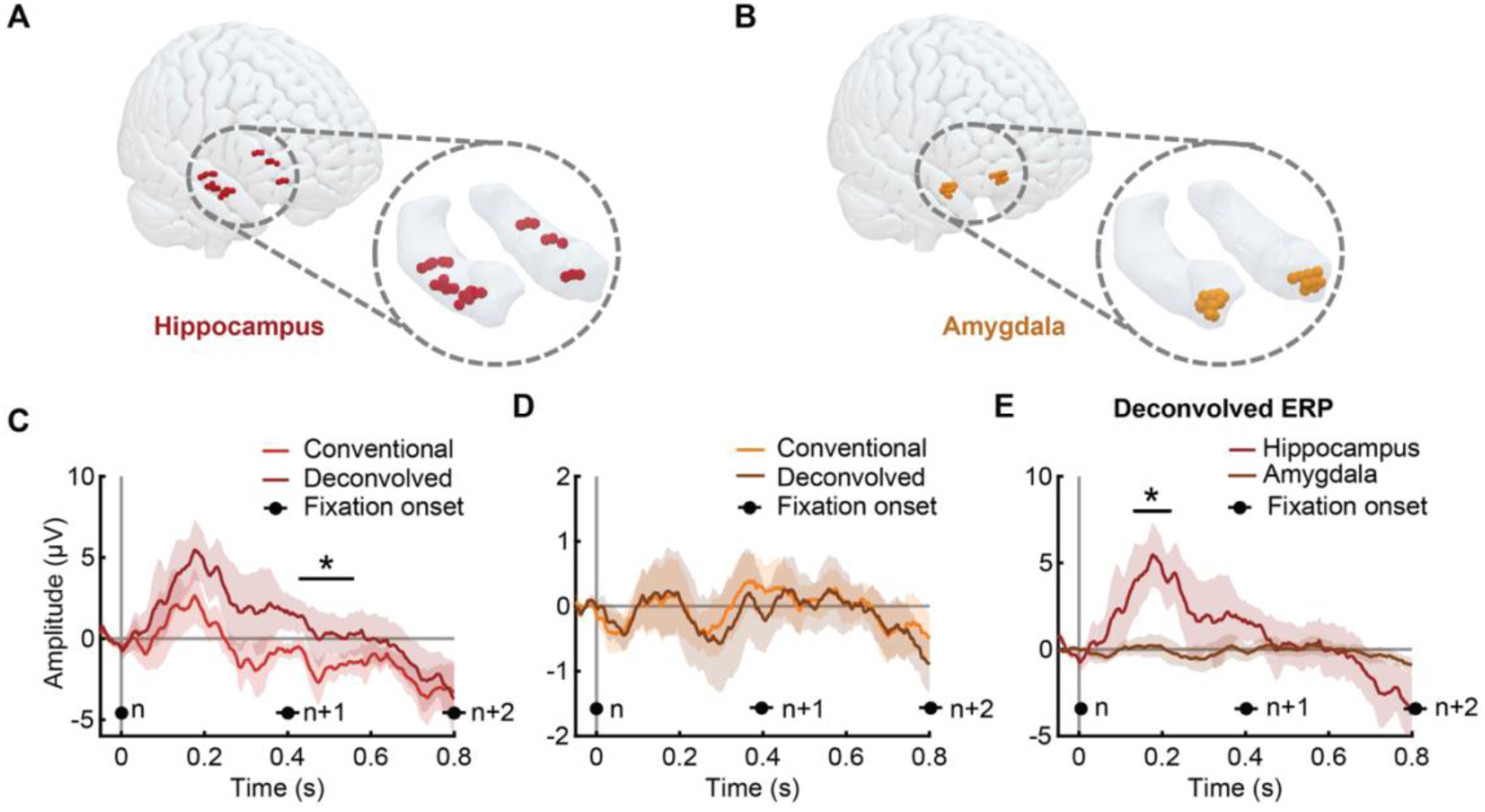
Deconvolutional modeling isolated fixation-evoked activity in the hippocampus. Intracranial electrode locations in hippocampus (maroon, **A**) and amygdala (orange, **B**). Conventional and deconvolved fixation-locked ERPs in the hippocampus (**C**) and amygdala (**D**). For ERPs, shaded regions denote SEM. Mean fixation onsets per fixation order (n, n+1, n+2) are indicated at the bottom of each panel, with lines denoting SEM. **E.** Direct comparison of deconvolved fixation-locked ERPs for hippocampus versus amygdala. Horizontal black lines indicate clusters with significant differences (* *p <* 0.05, corrected via permutation) for panels C and E.

The recording reference and ground of electrophysiological recordings consisted of either a surgically implanted electrode strip facing the scalp or a scalp electrode. Recordings were made using a Nihon Kohden amplifier with a sampling rate of 1–2 kHz, per clinical needs. Recorded signals were bandpass filtered from 0.6 to 600 Hz, re-referenced offline to a bipolar montage computed using adjacent electrode contacts and re-sampled to 1000 Hz.

### Experimental Design and Statistical Analysis

#### Behavioral and eye-movement analyses

We first assessed whether episodic memory performance and eye-movement behavior differed among semantic categories. Specifically, we tested whether (i) recognition accuracy and (ii) fixation counts varied by scene category; whether (iii) the number of fixations (both overall and On-Category fixations) predicted scene recognition; and whether (iv) the semantic category of the first fixation predicted recognition success.

For these behavioral tests, we fitted linear mixed-effects models with participant treated as a random effect. Fixed effects included semantic category and, where appropriate, fixation-based predictors. Estimated marginal means and pairwise contrasts between conditions were computed from fitted models. Multiple comparisons were corrected using the Benjamini–Hochberg false discovery rate (FDR) procedure (Models i and ii) or using parametric bootstrap tests (Models iii-iv). Multiple-comparisons correction strategies were chosen based on model complexity and the inferential goals of each analysis.

Model specifications were as follows:

i. Recognition accuracy

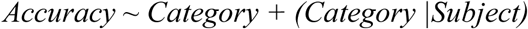

1. ii. Fixation count

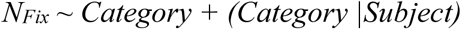

1. iii. Fixation count and recognition accuracy

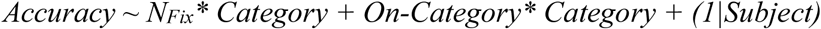

1. iv. First-fixation content and recognition accuracy

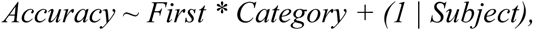

Where *Accuracy* is the recognition probability of hit during the test phase of the memory task; *Category* is the scene category (People, Animals, Food); *N_Fix_* is the number of fixations during a scene; *On-Category* specifies whether a fixation is made to the condition-defining content (i.e., either On-Person, On-Animal, On-Food or not); and *First* specifies whether the first fixation was made to the condition-defining content. For generalized LMEs (models iii and iv), *N_Fix_* and *On-Category* predictors were scaled to zero mean and unit variance within participant.

To test the specificity of first fixation effects, we included a control analysis contrasting effects with second and third fixations. We fitted a generalized linear mixed-effects model with fixed effects including content viewed (On/Off-Category), fixation position (First, Second, Third), and their interaction, while subject was included as a random intercept. Using the estimated marginal means of the model, we calculated the linear trend of the On-Category effects across fixation positions using linear contrast.

#### Deconvolution modeling of fixation-locked iEEG signals

Eye movements occur in rapid succession during scene viewing, potentially resulting in substantial temporal overlap of neural responses elicited by scene presentation, fixations, and saccades (Kragel and Voss, 2022). Therefore, fixation-locked neural activity can be confounded by overlapping activity elicited by subsequent fixations or to the initial scene onset itself. This overlap of neural signals complicates interpretation of the specific activity that eye movements and other task-related events trigger whenever eye movements occur (Kragel and Voss, 2022).

To address this challenge, we used a deconvolution approach using the Unfold toolbox for MATLAB (Ehinger & Dimigen, 2019). This method models the continuous electrophysiological signal as a linear combination of overlapping event-related responses, allowing fixation-evoked activity to be disentangled from scene-evoked activity and from other temporally adjacent fixations. We implemented two complementary deconvolution models. The first estimated a generalized fixation-locked ERP across all fixations, enabling direct comparison with conventional fixation-locked ERPs. The second estimated category- and order-specific fixation-locked ERPs, allowing us to examine how semantic content and temporal order of fixations modulated fixation-related activity.

Continuous iEEG data from all encoding blocks were concatenated for each participant. Scene-onset events were identified from task markers and labeled by the scene-level semantic content (People, Animals, Food). Fixation events were extracted from the eye-tracking parser and annotated according to the category of the fixated location. Then, they were labeled with their fixation order within the scene (i.e., first fixation, second fixation, etc.), and whether the fixated location was to the semantic content (On-Person, On-Animal, and On-Food fixations) or to other locations of the scene that did not have the scene-specific semantic content (Off-Person, Off-Animal, and Off-Food).

We fitted two different deconvolution models matching our goals. In each model, we fitted neural responses to both the scene onset and fixations, which we specify here expressed in Wilkinson notation {scene, fixation}. The first general model estimated responses to all scene onset and fixation events:

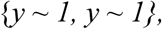

where *y* is the continuous encoding period data from either the amygdala or hippocampus. To examine semantic and order effects, we specified a second model:

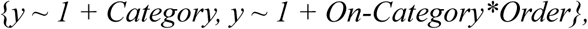

where *y* is the continuous iEEG data, *Category* is the semantic category of the scene, *On-Category* specifies the semantic content of a fixation, and *Order* is the temporal order of fixations in the viewing sequence (grouped as either the first fixation or fixations later than the first). For both models, regressors were temporally expanded from –100 to 800 ms around each event. Epileptiform activity during these latencies was detected using a kurtosis threshold of 10 and trials with this pathological activity were excluded. The deconvolution model was then fitted to the continuous signal, yielding time-resolved beta estimates that correspond to deconvolved scene-evoked and fixation-evoked responses for each electrode, semantic content, and temporal order of fixations. Then, we averaged beta estimates across electrodes for each participant and brain region of interest (hippocampus and amygdala).

We also computed conventional fixation-locked ERPs by segmenting raw iEEG data into epochs time-locked to fixation onset (–100 ms to 800 ms) and averaging across trials. We discarded trials with epileptiform activity as well, using the kurtosis threshold described above. Artifact-free epochs were averaged, first within electrodes and then across electrodes for each participant and brain region of interest.

Before group-level analyses, conventional fixation-locked ERPs and deconvolved ERPs were baseline-corrected using a −50 to 0 ms pre-fixation window and line-noise corrected. Group-average waveforms were obtained by averaging across participants, with inter-subject variability quantified using the standard error of the mean (SEM).

#### Statistical analysis of conventional and deconvolved fixation-locked ERPs

Conventional and deconvolved fixation-locked ERPs were statistically compared to evaluate the impact of deconvolution on the estimated neural responses. We performed a within-subject cluster-based permutation test using FieldTrip toolbox comparing conventional and deconvolved fixation-locked ERPs in hippocampus and amygdala separately (Maris and Oostenveld, 2007). For each participant, we performed dependent-samples t-test using a two-tailed cluster correction with 10,000 randomizations. The test returned latency-interval clusters with significant amplitude differences for conventional versus deconvolved responses, providing an assessment of how the deconvolved activity differed from the conventional ERP in estimation of fixation-related signal. To compare the deconvolved ERPs between brain regions (hippocampus and amygdala) and thus, assess the regional specificity of the fixation-evoked effects, we used the same cluster-based permutation test constrained to the latencies around the evoked response (100 to 300 ms after the fixation onset).

Deconvolved fixation-locked ERPs were compared across multiple conditions to examine semantic selectivity and fixation-order effects. We compared deconvolved responses among five conditions: (i) scene-locked ERPs, (ii) overall On-Category fixation-locked ERPs (iii) first On-Category fixation-locked ERPs and (iv) later (second and onward) On-Category fixation-locked ERPs, compared among semantic categories, as well as (v) overall fixation-locked ERPs compared between temporal order categories (first versus above).

For each comparison, we performed a repeated-measures ANOVA with three levels corresponding to the three semantic categories. Each ANOVA identified latency intervals with significant amplitude differences among categories after controlling for multiple comparisons via cluster permutation (10,000 iterations). Post hoc analyses were conducted following the significant ANOVA across semantic fixation categories. Pairwise comparisons between conditions (On-person vs On-animal, On-person vs On-food, and On-animal vs On-food) were performed using cluster-based permutation tests. To control for multiple comparisons across post hoc clusters, p-values were further corrected using the FDR procedure (q = 0.05).

We also tested brain-region specificity for the first-fixation effects, comparing first On-Person fixation-locked ERPs between hippocampus and amygdala following the same statistical methodology described above to compare brain regions.

For each significant comparison, we computed Hedges’ g for dependent samples on the average response within the significant cluster to quantify the magnitude of the effect.

#### Theta phase-locking computation and statistical analyses

Theta phase-locking analyses characterized fixation-related neural frequency-domain dynamics. Human hippocampal theta-band oscillatory activity (∼3 – 8 Hz) is associated with episodic memory encoding, exhibiting systematic phase modulation during successful memory formation (Lega et al., 2012; Cruzat et al., 2021). Eye movements are coupled with phase resetting of theta oscillations, which during encoding predicts successful episodic memory formation (Jutras et al., 2013; Kragel et al., 2020). Accordingly, we analyzed theta phase locking changes relative to fixations to capture theta-phase mechanisms supporting episodic memory encoding, complementary to the evoked-potentials analyses described above.

For this analysis, continuous recordings were segmented into trials aligned to scene onset. These trials were epoched in latency intervals of 0.8 to 4 s relative to scene onset to avoid edge artifacts from wavelet convolution. For detection of oscillatory peaks, power spectra were first computed using multitaper Fourier analysis with a Hanning taper (1–40 Hz, FFT-based). Power spectra were log-transformed and averaged across trials. To isolate oscillatory components, the aperiodic (1/f) component of the spectrum was estimated using a robust fitting procedure (Kragel et al., 2020) and subtracted from the power spectrum, yielding a whitened spectrum. The dominant oscillatory peak frequency was identified for each electrode as the maximum power within the 2.5–10 Hz range of the whitened spectrum, corresponding to the approximate potential theta band. This peak frequency was then used for all subsequent phase analyses for that electrode.

Time-resolved Fourier coefficients were computed at the electrode-specific peak frequency using a multitaper convolution with a window length of five cycles. Instantaneous phase and amplitude were extracted from the complex coefficients. Phase values were then realigned to fixation onset by extracting a −100 ms to +100 ms latency interval centered on each fixation. Analyses were conducted either based on fixation order (first, second, etc.) or by collapsing across all fixations. Fixations were grouped by semantic content (On-Person, On-Animal, On-Food, or Off-Category).

Phase consistency across fixations was quantified using the mean resultant vector length (r), computed separately for each semantic content, time point, and fixation order. To assess statistical significance, null distributions were generated via permutation testing. For each electrode, fixation labels were randomly shuffled 1,000 times prior to fixation alignment, thereby disrupting the relationship between fixation timing and neural phase while preserving the temporal and spectral structure of the data. Phase consistency metrics were recomputed for each permutation, yielding null distributions for each semantic category, time point and fixation order.

Observed phase-locking values were Fisher-z–transformed, averaged across participants, and compared against their corresponding null distributions using permutation tests. Two-sided p-values were computed as the proportion of null statistics exceeding the observed test statistic. Bootstrap confidence intervals (1,000 iterations) were computed for group-level phase-locking estimates.

To assess the specificity of first fixation effects on the semantic modulation of theta-phase locking, we analyzed the linear trend of phase-locking values of the first three fixations in hippocampus and amygdala. For each subject, phase-locking values and their corresponding null distributions related to On-Person fixations were corrected by subtracting the mean phase-locking value across fixations and computing a weighted average (decreasing linearly with fixation order) across fixations. Group-level statistical significance was assessed using permutation testing, comparing observed responses against the null distribution.

## Results

### Semantic content and fixation order jointly predict recognition for natural scenes

We tested whether scenes with different semantic content (People, Animals, and Food) differed in recognition accuracy or viewing behavior during encoding. Recognition accuracy did not differ based on semantic content (proportion correct for People: 0.70 ±0.06, Animals: 0.80 ± 0.04, Food: 0.76 ± 0.04), nor did the mean number of fixations during encoding (People: 6.70 ± 0.31, Animals: 6.55 ± 0.44, Food: 7.24 ± 0.58; Figure 1B). However, the number of fixations to On-Category locations during encoding predicted later recognition. We modeled recognition accuracy as a function of the total number of fixations and the number of fixations to On-Category locations (i.e., On-Person, On-Animal, On-Food; Figure 1A). More fixations to On-Category locations correlated significantly with higher recognition accuracy (χ²(1) = 7.89, p = 0.011; Figure 1C), whereas the total number of fixations overall did not significantly correlate with recognition accuracy (χ²(1) = 0.63, p = 0.52). This correlation did not significantly differ among semantic categories (χ²(2) = 2.22, p = 0.34) and there was no significant interaction of fixation number with semantic category (On-Category fixations: χ²(2) = 1.75, p = 0.42; total fixations: χ²(2) = 3.49, p = 0.17). Thus, more fixations to semantic content within scenes predicted better recognition memory, without significant variation among the three semantic categories.

We next tested the hypothesis that fixation order influences semantic processing relevant to memory formation. For People scenes, subsequent recognition was significantly higher when the first fixation during encoding was centered on person content than when first fixations landed on non-person semantic content within the scenes (On-Person versus Off-Person; odds ratio = 0.48, z = 2.44, p = 0.015; Figure 1D left). No corresponding effects were observed for Animal scenes (On-Animal vs. Off-Animal; odds ratio = 1.031, p = 0.93; Figure 1D middle) or Food scenes (On-Food vs. Off-Food; odds ratio = 0.69, p = 0.24; Figure 1D right). We did not observe similar effects for second or third fixations on People scenes (On-Person vs. Off-Person; second fixation: odds ratio = 0.68, p = 0.24; third fixation: odds ratio = 1.21, p = 0.55). For People scenes, there was a significant linear decrease for subsequent memory across first, second, and third fixations (z = 2.12, p = 0.034), indicating that the difference in subsequent recognition for On-Person versus Off-Person fixations decreased systematically from the first to later fixations. We did not observe similar changes for Animal (linear: z = 0.25, p = 0.80) or Food (linear: z = −0.52, p = 0.60) scenes.

These results indicate that, whereas overall recognition accuracy and the number of fixations were matched among scene categories, memory varied according to whether people features were rapidly fixated, on the first versus subsequent fixations, in People scenes. This supports the hypothesis that first fixations play a privileged role in memory formation and that this role is selective, depending on semantic content.

### Isolation of fixation-evoked iEEG activity in hippocampus

We used deconvolutional modeling to isolate iEEG activity evoked by specific fixations from potentially overlapping activity associated with the overall scene onset and with activity evoked by other, temporally proximal fixations. This analysis approach was essential because fixations occurred within hundreds of milliseconds after scene image onset, during the time when neural processing of the scene would be expected to occur (Puce et al., 1997; Paller and McCarthy, 2002). Moreover, field potentials evoked by fixations or saccades last for hundreds of milliseconds (Katz et al., 2020). Because fixations occurred in rapid succession (one fixation every 400 ± 30 ms), there was substantial potential for temporal overlap of the neural responses to scenes and to consecutive fixations. To address this, we used deconvolution modeling to isolate activity uniquely associated with each individual fixation, taking scene onsets and other fixations into account. To validate that deconvolutional modeling isolated iEEG signals of fixations, we compared the fixation-locked deconvolved ERPs to those quantified via conventional fixation-locked methods, without any deconvolutional modeling, in which case activity related to scene and fixations onsets are not separated and can jointly influence activity in unknown proportions.

In hippocampus, conventional fixation-locked ERPs showed a multiphasic response with relatively low amplitude (Figure 2C). In contrast, deconvolved ERPs showed a larger evoked response with a single, clear peak, without multiphasic activity (Figure 2C). Cluster-based permutation testing revealed one significant latency interval during which deconvolved ERPs were significantly larger than conventional ERPs (444 to 542 ms post fixation onset; p = 0.029; g = 0.65, 95% CI [−0.01, 1.46]). To aid in interpreting the timing of this effect, we show the average onset times of three fixations: the primary fixation at time zero of the ERP, and the following two fixations (plotted above the x-axis in Figure 2C,D). We observed significant differences between conventional and deconvolved ERPs following the average onset of the second fixation, indicating that conventional ERPs substantially underestimate hippocampal responses due to interference from overlapping activity from other fixations and/or scene onsets. Thus, deconvolutional modeling was important for isolating fixation-evoked neural activity in hippocampus.

There were no robust fixation-locked ERPs for amygdala and no effect of deconvolutional modeling on ERP amplitude for amygdala (Figure 2D). Deconvolved fixation responses were significantly greater in hippocampus than amygdala within an early latency interval (147–205 ms; p = 0.046, g = 0.31, 95% CI [−0.98, 1.69]; Figure 2E). Thus, fixation-locked ERPs were specific to hippocampus. In addition to suggesting functional differences between these structures, this finding is important in ruling out a major potential confounding factor in the analysis of fixation-locked ERPs. Movement of the corneo-retinal dipole can induce electrical signals (Plöchl et al., 2012), as can contraction of extraocular muscles involved in saccadic eye movements (Berg and Scherg, 1991). The amygdala is closer to the orbit than hippocampus, making amygdala electrodes more likely to capture any purely movement-related electrical activity, which would be considered noise for our analysis goals. Thus, these findings indicate that fixation-locked activity of hippocampus was likely of neural origin.

### Semantic and temporal selectivity of fixation-evoked hippocampal activity

As the first step towards identifying hippocampal neural correlates of the behavioral finding that first On-Person fixations during scene encoding predicted subsequent recognition (Figure 1D), we tested whether fixation-locked hippocampal activity was sensitive to semantic content. Indeed, the amplitude of deconvolved fixation-locked ERPs in the hippocampus varied with the semantic content of On-Category fixations. Cluster-based permutation ANOVA indicated significant differences among semantic categories at 292-502 ms post-fixation (p = 0.006; Figure 3A). Post hoc pairwise comparisons indicated the most positive ERP amplitudes for On-Person fixations (average in cluster: 9 ± 5 µV), compared to On-Animal fixations (−1 ± 4 µV; p < 0.001, g = 0.86, 95% CI [0.099, 1.83]) and On-Food fixations (3 ± 5 µV; p < 0.001, g = 0.40, 95% CI [-0.09, 0.99]).

**Figure 3.**
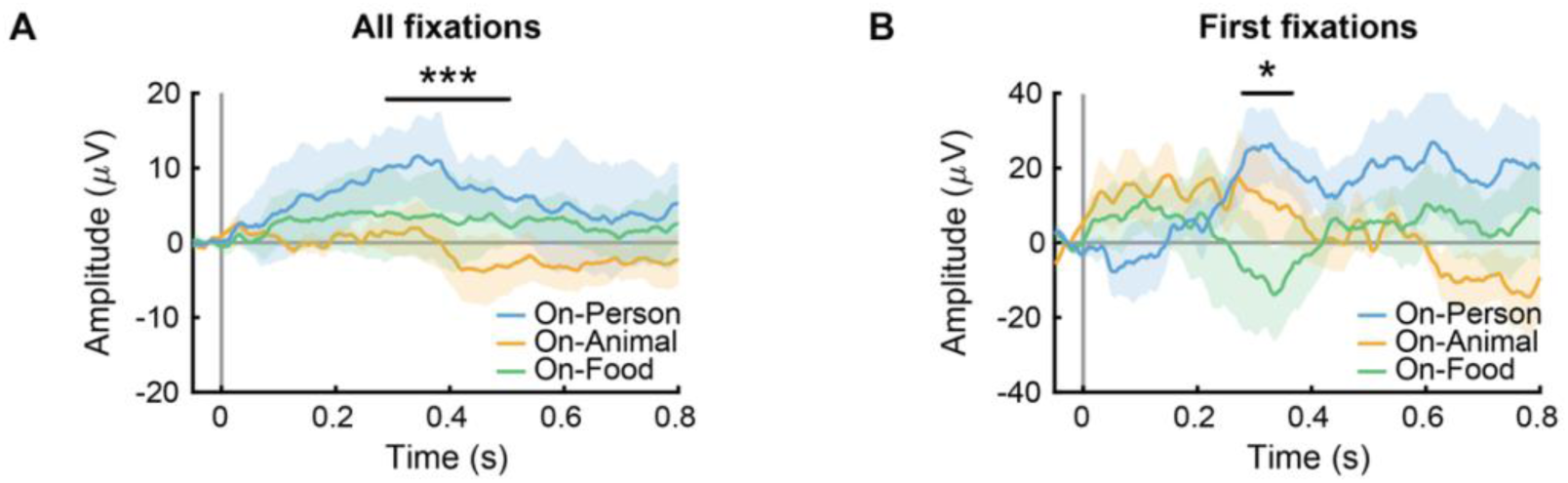
Content-selectivity of hippocampal activity occurs as early as the first fixation. **A.** Deconvolved fixation-locked ERPs for all On-Category fixations (On-Person, On-Animal, On-Food). **B.** The same semantic categories for only the first fixations during study trials. Shaded regions denote SEM. Horizontal black lines indicate clusters with significant between-category differences (* *p <* 0.05, *** *p < 0.001*, corrected via cluster permutation).

In contrast, scene-locked ERPs (defined based on the onset of the visual scenes rather than the onset of fixations) did not differ among semantic categories (no significant clusters detected by the ANOVA testing differences among categories). Thus, the effects of semantic category on hippocampal iEEG responses were specific to fixations.

We next assessed whether the ERP differences among categories were sensitive to the temporal order of fixations. Thus, we compared deconvolved ERPs among semantic categories for the first On-Category fixations and for subsequent On-Category fixations (i.e., the second and later fixations). The effects of semantic content varied by temporal order. That is, ERP amplitude for On-Category fixations varied by semantic category for the first fixation during encoding trials (281-365 ms; p = 0.047; Figure 3B). ERP amplitudes during this latency interval were more positive for On-Person fixations (24 ± 10 µV) than for On-Animal fixations (11 ± 11µV) and On-Food fixations (−10 ± 13 µV; Figure 3B). Post-hoc analysis revealed significantly higher amplitudes in the first On-Person fixation ERP compared to first On-Food fixation ERP (p = 0.031) that did not survive FDR correction (q = 0.063). Further, there were no significant effects of semantic content for On-Category fixations after the first fixation during encoding (i.e., second fixation and above; no significant clusters were identified). Thus, hippocampal fixation-locked activity was influenced by semantic content and fixation order in a manner that paralleled the behavioral findings. That is, first fixations to People semantic content were relevant for subsequent memory (Figure 1D) and for fixation-locked hippocampal activity (Figure 3B).

To test an alternative possibility that there was a general effect of order irrespective of semantic category, we compared between first and above fixation-locked ERPs without separating by semantic categories. There were no significant effects of temporal order (first vs. later fixations) on hippocampal fixation-locked deconvolved ERPs in general, pooling across semantic categories (no significant clusters detected).

We next tested specificity to hippocampus by performing the same set of analyses for amygdala. For amygdala, there were no significant effects of semantic category for any comparison of fixation-locked deconvolved ERPs (no significant clusters detected). To test whether the positive ERP amplitudes identified for first On-Person fixations in hippocampus (Figure 3B) were selective, we compared ERPs for this condition in hippocampus versus amygdala. Amplitudes were significantly greater in hippocampus relative to amygdala from 226 – 420 ms (hippocampus 19 ± 0 µV; amygdala −1 ± 0 µV; p = 0.046, g = 1.27, 95% CI [-0.107, 2.952]). Notably, this latency interval of significant hippocampus versus amygdala difference identified via permutation testing coincided with the latency interval of the effect of semantic category for first fixations in hippocampus (Figure 3B). These results indicate that semantic selectivity in evoked responses to first fixations was specific to hippocampus.

### People-selective theta phase locking emerges during the first fixation

To characterize fixation-related oscillatory dynamics, we examined theta phase locking to fixation onsets in hippocampus and amygdala. Clear oscillatory peaks were apparent after removing the aperiodic component of the power spectrum in the low theta (2.5–5 Hz) and high theta (5–10 Hz) bands in both the hippocampus (Figure 4A) and amygdala (Figure 4B). For each electrode, we then computed phase locking at the dominant theta frequency during fixations to each semantic category, comparing observed phase-locking values to a null distribution generated by shuffling fixation labels (see Methods).

**Figure 4.**
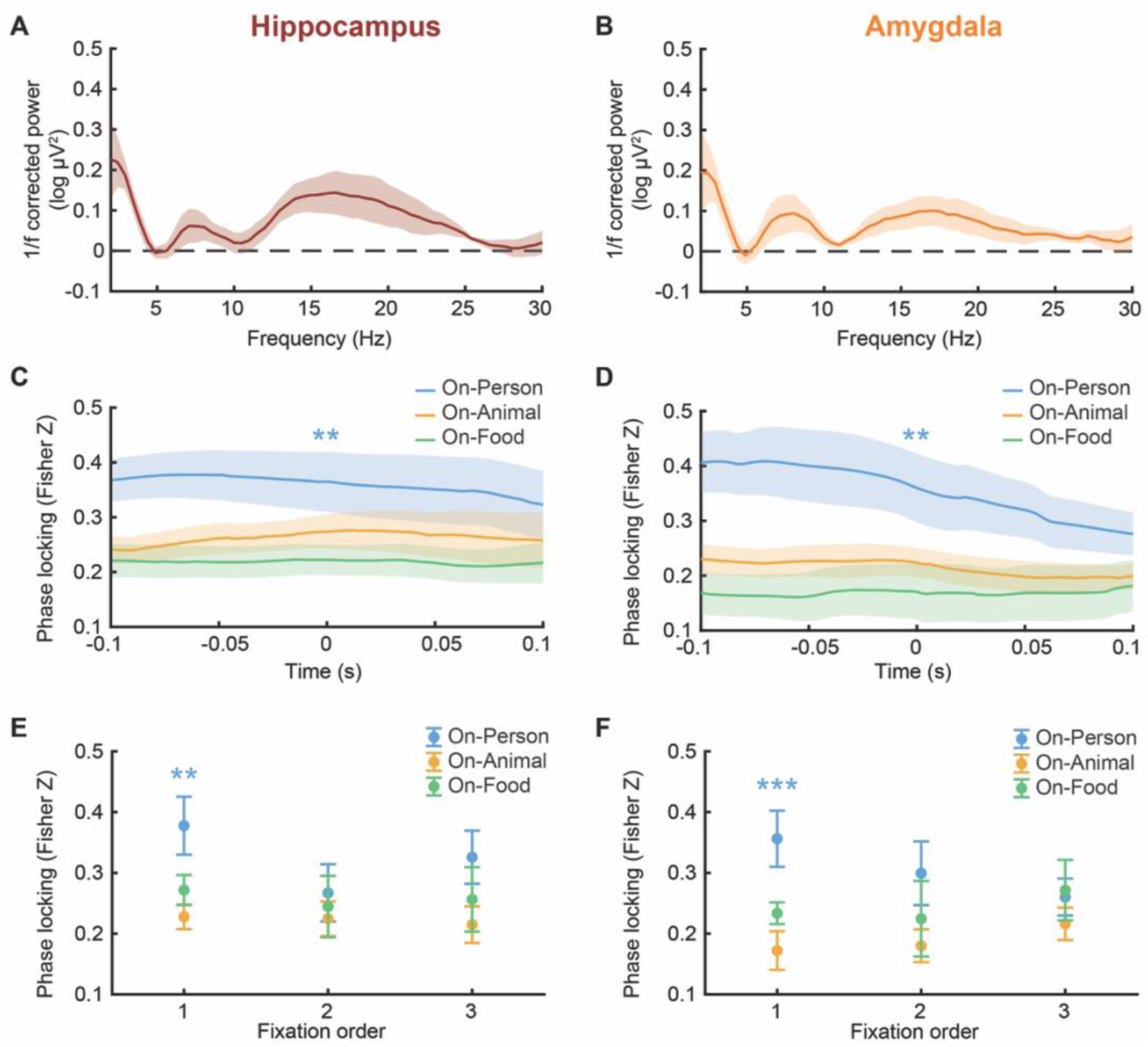
Theta phase locking to fixations varies by semantic category and fixation order in hippocampus and amygdala. **A** and **B.** Theta oscillatory peaks in hippocampus (A) and amygdala (B). **C** and **D.** Comparison of theta phase locking to fixations among semantic categories for hippocampus (C) and amygdala (D). **E** and **F.** Comparison of theta phase locking to fixations among semantic categories and early-trial fixation order for hippocampus (E) and amygdala (F). ** p < 0.01. *** p < 0.001.

We first tested whether phase locking varied by semantic category. In hippocampus, On-Person fixations had significantly stronger theta phase locking than expected by the null distribution (Δz = 0.19, p = 0.009; Figure 4C), whereas On-Animal fixations (Δz = 0.05, p = 0.84) and On-Food (Δz = 0.09, p = 0.41) did not. There were similar findings in amygdala, which had significant theta phase locking for On-Person fixations relative to the null distribution (Δz = 0.21, p = 0.003), whereas On-Animal fixations (Δz = 0.02, p = 0.98) and On-Food fixations (Δz = 0.07, p = 0.77; Figure 4D) did not.

The selective theta phase locking identified for On-Person fixations was specific to first fixations during encoding trials. Theta phase locking significantly exceeded the null distribution for first On-Person fixations in both hippocampus (Δz = 0.36, p = 0.002; Figure 4E) and amygdala (Δz = 0.34, p < 0.001; Figure 4F). No significant phase locking was observed for second (Δz = 0.26, p = 0.34) or third (Δz = 0.31, p = 0.098) On-Person fixations in hippocampus or amygdala (second: Δz = 0.29, p = 0.076; third: Δz = 0.26, p = 0.69). Similarly, there were no significant effects for first or later On-Animal (first fixation: Δz = 0.22, p = 0.42; second fixation: Δz = 0.22, p = 0.34; third fixation: Δz = 0.21, p = 0.90) or On-Food fixations (first fixation: Δz = 0.26, p = 0.098; second fixation: Δz = 0.24, p = 0.17; third fixation: Δz = 0.25, p = 0.21) in hippocampus. Likewise, there were no significant phase-locking effects in amygdala for On-Animal fixations (first fixation: Δz = 0.17, p = 0.99; second fixation: Δz = 0.18, p = 0.93; third fixation: Δz = 0.21, p = 0.90) or On-Food fixations (first fixation: Δz = 0.23, p = 0.46; second fixation: Δz = 0.22, p = 0.39; third fixation: Δz = 0.27, p = 0.057).

To quantify decreases in phase locking later in the fixation sequence (Fig. 4E,F), we computed a linear contrast across the first three fixations (see Methods) and compared these measures to a permutation null, shuffling fixation labels across trials. Phase locking in hippocampus decreased across the first three fixations significantly more than the permutation null (z = 0.047 ± 0.038, p = 0.026). In contrast, there was no significant effect of fixation order in amygdala (z = 0.006 ± 0.041, p = 0.50), where the observed response did not differ from the null distribution. These findings indicate that viewing people selectively modulated hippocampal and amygdala theta dynamics at the earliest moments of visual exploration.

### Control analyses of saccade and fixation durations

To test the possibility that observed neural differences may have reflected eye-movement properties rather than stimulus-related neural processing per se, we compared fixation and saccade durations for the conditions that were associated with effects on deconvolved fixation-locked ERP amplitudes in the aforementioned analyses.

On-Category fixation durations did not vary significantly among semantic categories (On-Person fixations 405 ± 28 ms; On-Animal fixations 429 ± 36 ms; On-Food fixations 363 ± 34 ms; F(2, 7.01) = 3.59, p = 0.085). Likewise, fixation durations did not vary among semantic categories when analyzing only the first fixations (On-Person 403 ± 38 ms; On-Animal 353 ± 33 ms; and On-Food 329 ± 42 ms; F(2, 7.59) = 1.31, p = 0.32).

On-Category saccade durations also did not vary significantly among semantic categories (On-Person 33 ± 2 ms; On-Animal 32 ± 3 ms; On-Food 34 ± 2 ms; F(2, 5.70) = 1.34, p = 0.33). Likewise, saccade durations did not vary among semantic categories when analyzing only the first fixations (On-Person 29 ± 2 ms; On-Animal 27.84 ± 1 ms; On-Food 28 ± 2 ms; F(2, 7.69) = 0.36, p = 0.71). These findings indicate that effects of semantic category and fixation order on fixation-locked neural signals were unlikely to have resulted indirectly from corresponding differences in fixation or saccade properties.

## Discussion

By isolating neural responses to eye fixations using a deconvolution-modeling approach, the analyses reported here establish that hippocampal iEEG activity during scene encoding is organized around discrete fixation events and is sensitive to their semantic content and temporal order. Specifically, fixations to people modulate fixation-locked hippocampal responses, expressed as both larger fixation-evoked potentials and stronger theta phase locking relative to fixations on animals or food. Moreover, this semantic sensitivity was driven by the first fixation, relative to later fixations within a trial. Subsequent recognition memory performance varied with whether the first fixation was directed to people versus to other content within the same scenes. Together, these findings suggest that for scenes of people the first fixation acts as a temporally crucial moment for hippocampal memory encoding.

One major implication of these findings is that fixational eye movements provide temporal structure to hippocampal semantic processing. Prior work has demonstrated that hippocampal neurons respond selectively to specific categories of visual stimuli, individual objects, and familiar people (Quiroga et al., 2005; Quiroga, 2012; Mackay et al., 2024; Bausch et al., 2026), but has largely not assessed the temporal relationship between such selectivity and discrete sampling events. Although semantic information could theoretically be processed continuously during stimulus viewing, without regard to the timing of eye movements, the current findings suggest otherwise. Our deconvolution approach allowed separation of stimulus-evoked and fixation-evoked iEEG activity, and we found no significant semantic-related variation in scene-locked hippocampal responses—only in fixation-locked responses. This suggests that hippocampal sensitivity to semantic content is synchronized to fixations at least for the first fixations to people. This is analogous to how, during reading, viewing of each word triggers neural signals of semantic processing (i.e., N400 ERPs) (Federmeier, 2022). Notably, the latency at which fixation-locked hippocampal ERPs were sensitive to semantic content (∼400 ms) is similar to N400 latencies observed in reading and other contexts (Federmeier, 2022), including to N400s measured to specific fixations isolated via the deconvolution method used here (Coco et al., 2020). This consistency raises the possibility that incorporation of stimulus meaning in hippocampal memory processing occurs discretely during specific latencies relative to fixations, following the same general conceptual processing in non-hippocampal portions of the inferior and anterior temporal lobes (Nobre and McCarthy, 1994, 1995; McCarthy et al., 1995).

Theta phase locking to fixation onset provides a plausible mechanism for this discretized memory processing. Phase resetting is thought to create time-limited windows during which hippocampal circuits can effectively encode incoming information (Buzsáki, 2002; Sauseng et al., 2007; Lisman and Jensen, 2013) including during visual exploration (Jutras et al., 2013; Kragel et al., 2020; Staudigl et al., 2022). We observed that phase locking of theta was selectively enhanced for fixations to people, which was the same condition that drove semantic sensitivity in the evoked response. Such co-occurring enhanced evoked responses and theta phase locking under the same conditions suggests these are complementary rather than redundant measures. Fixation-locked hippocampal responses may index hippocampal engagement with fixation-locked phase alignment establishing a temporal window within which that engagement is coordinated across the medial temporal lobe network.

The dissociation between hippocampus and amygdala across the two measures is informative. Whereas fixation-evoked amplitude enhancement was specific to hippocampus, theta phase locking was observed in both hippocampus and amygdala. This pattern is consistent with distinct but complementary roles for the hippocampus and amygdala during scene viewing. That is, the hippocampus might encode relational and contextual structure at fixations, whereas amygdala might signal the behavioral relevance of fixated content. That both effects were driven specifically by fixations to people, a category of innate biological value for which amygdala orienting is selectively engaged (Kaskan et al., 2022), suggests these responses reflect coordinated but functionally distinct processes.

Our scenes depicted people embedded in rich social and spatial environments, engaging in activities, interacting with objects, and occupying recognizable settings. This experimental context is qualitatively different from the isolated faces or artificial arrays used in typical amygdala face-processing studies (Rutishauser et al., 2011; Wang et al., 2014; Becker and Rheem, 2020). In such naturalistic contexts, fixating a person involves sampling a structured event whose meaning depends on relational context, which may preferentially engage hippocampus over amygdala.

Although fixating a face in a naturalistic scene can evoke prototypical signals of face processing (Auerbach-Asch et al., 2020; Gert et al., 2022), the contribution of the amygdala here may be better characterized as relevance signaling that orients the broader medial temporal lobe network toward the fixated content, rather than semantic encoding per se.

Theta phase locking in both amygdala and hippocampus is consistent with this view. The amygdala may participate in a network-level phase resetting response driven by the relevance of semantic content, signaled or coordinated by the hippocampus without independently generating the encoding-related amplitude modulation—consistent with its role in presaccadic orienting to biologically significant targets (Peck et al., 2013; Lee et al., 2026). Such coordination likely involves multiple amygdalar nuclei and subcortical pathways, and future studies with simultaneous recordings across these structures would be needed to confirm such a role (Janak and Tye, 2015; Maeda et al., 2020). Inter-regional theta dynamics between the amygdala and hippocampus (Zheng et al., 2017; Staudigl et al., 2022) further support coordinated processing, though determining directionality would necessitate simultaneous recordings in upstream regions such as the medial septum or entorhinal cortex (Buzsáki, 2002; Colgin, 2016) to rule out a common input driving both structures.

The concentration of both neural effects, fixation-evoked amplitude and theta phase locking, during the first fixation deserves emphasis. First fixations may serve not as typical sampling events but as anchoring moments that shape how subsequent scene information is interpreted and encoded. When the first fixation falls on a person, it may instantiate a contextual or relational reference in which the rest of the scene is understood. What the person is doing, who they are interacting with, or what their actions may signify impart meaning to the scene. In contrast, types of semantic content such as animals or food may not require referential context for relational processing given the contexts in which they typically occur or the actions or interactions they naturally afford.

The memory benefit of first fixations to people may arise not simply from the encoding of that fixation itself, but from the downstream consequences of establishing a visual context early in the viewing sequence. This interpretation is consistent with the role of the hippocampus in building and exploiting relational and contextual structure (Eichenbaum, 2004, 2017; Konkel and Cohen, 2009), with evidence from scalp EEG that fixation-locked theta activity predicts memory (Nikolaev et al., 2023), and with evidence that memory for scenes is enhanced when viewers construct coherent event representations during encoding (Zacks et al., 2007).

Indeed, previous studies have demonstrated that the first fixation on a scene differs from subsequent fixations (Arizpe et al., 2012; Minxha et al., 2017), often reflecting rapid extraction of global or semantically salient information (Loftus and Mackworth, 1978; De Graef et al., 1990; Castelhano and Henderson, 2003) that is relevant to memory formation (Loftus and Mackworth, 1978; Wynn et al., 2016). We thus suggest that first fixations to people may have influenced hippocampal activity and subsequent memory not because the stimuli were social or salient per se, but because they provided a context for interpreting the relations among subsequently viewed information in the scene. This is broadly consistent with the established role of fixational eye movements in relational aspects of hippocampal-dependent memory (Hannula et al., 2007; Hannula and Ranganath, 2009). Our results thus provide novel support for a model whereby hippocampal engagement during visual exploration is gated by the semantic content of information being sampled, particularly when that content carries rich relational and contextual structure.

Several limitations should be considered. First, the sample of participants was small (n = 6) and comprised patients with medically refractory epilepsy, which may limit the generalizability of the results to healthy populations, as underlying pathology could influence brain activity. To mitigate this concern, trials containing epileptiform activity were identified and excluded from all analyses. Second, we investigated only three stimulus categories, which constrains comparisons with broader categorical distinctions and may limit conclusions about category-general mechanisms. Future work should extend these analyses to larger and more expansive stimulus sets, including stimuli that more systematically vary the complexity and relational content of scenes. Finally, while our results establish a strong association between fixation timing, hippocampal dynamics, and memory, causal relationships remain to be determined.

In summary, our study demonstrates that both the semantic content and temporal order of visual fixations shape hippocampal activity and subsequent episodic memory. Specifically, we show that the first fixation on a person elicits the strongest fixation-locked evoked response in the hippocampus and theta phase locking in the hippocampus and amygdala, highlighting a unique neural mechanism by which gaze behavior modulates memory encoding. These findings establish that fixation-related hippocampal activity is not a uniform response to eye movements but is selectively modulated by what is being fixated and when. In doing so, they point toward a model in which theta-paced visual processing organizes the transformation of semantic and relational scene content into episodic memory.

